# Integrated analysis of single-cell embryo data yields a unified transcriptome signature for the human preimplantation epiblast

**DOI:** 10.1101/222760

**Authors:** Giuliano G Stirparo, Thorsten Boroviak, Ge Guo, Jennifer Nichols, Austin Smith, Paul Bertone

## Abstract

Single-cell profiling techniques create opportunities to delineate cell fate progression in mammalian development. Recent studies provide transcriptome data from human preimplantation embryos, in total comprising nearly 2000 individual cells. Interpretation of these data is confounded by biological factors such as variable embryo staging and cell-type ambiguity, as well as technical challenges in the collective analysis of datasets produced with different sample preparation and sequencing protocols. Here we address these issues to assemble a complete gene expression time course spanning human preimplantation embryogenesis. We identify key transcriptional features over developmental time and elucidate lineage-specific regulatory networks. We resolve post hoc cell-type assignment in the blastocyst, and define robust transcriptional prototypes that capture epiblast and primitive endoderm lineages. Examination of human pluripotent stem cell transcriptomes in this framework identifies culture conditions that sustain a naïve state pertaining to the inner cell mass. Our approach thus clarifies understanding both of lineage segregation in the early human embryo and of in vitro stem cell identity, and provides an analytical resource for comparative molecular embryology.

## Introduction

Three distinct cell lineages are established in the mammalian blastocyst: trophectoderm (TE) to support uterine implantation and development of the placental epithelia, extraembryonic primitive endoderm (PrE) from which the primary yolk sac is formed, and pluripotent epiblast (EPI) which gives rise to the embryo proper. In the human embryo, the first lineage segregation becomes apparent at embryonic days (E) 4–5, when the compacted morula undergoes cavitation to initiate blastocyst formation. Inner cells are directed towards the inner cell mass (ICM), while outer cells acquire TE fate.

Unlike in mouse, the pluripotency factor POU5F1 (also known as OCT4) is widely expressed in both early ICM and TE in the early human blastocyst (Niakan and Eggan, 2013). By E6 in human, POU5F1 is downregulated in TE but remains expressed in all cells of the ICM (Chen et al., 2009; Deglincerti et al., 2016; Niakan and Eggan, 2013). ICM cells with high POU5F1 levels often co-express NANOG (Roode et al., 2012; Deglincerti et al., 2016), suggesting a prospective EPI fate. Lower POU5F1 levels correlate with PrE-specific SOX17 expression (Roode et al., 2012; Niakan and Eggan, 2013).

Initial co-expression of NANOG and GATA6 in a subset of early ICM has also been observed in human (Roode et al., 2012) and non-human primates (Boroviak et al., 2015). In human blastocysts exceeding 200 cells, EPI and PrE marker profiles appear mutually exclusive (Niakan and Eggan, 2013): EPI cells are associated with NANOG and high POU5F1 expression, whereas PrE cells are characterised by GATA6, SOX17, GATA4 and diminishing levels of POU5F1 (Roode et al., 2012; Deglincerti et al., 2016). These features indicate EPI and PrE segregation in the peri-implantation blastocyst between E6 and E7. Selective in situ analyses have contributed seminal knowledge of key regulatory events that underlie early lineage progression in primate development. However, detailed characterisation of human embryogenesis on a genome-wide molecular level has been lacking.

Various high-throughput profiling methods have recently been applied to gene expression and DNA methylation analysis of embryos from several mammalian species, including mouse (Guo et al., 2010; Ohnishi et al., 2014; Boroviak et al., 2015; Guo et al., 2014), human (Xue et al., 2013; Yan et al., 2013; Blakeley et al., 2015; Petropoulos et al., 2016) and non-human primates (Boroviak et al., 2015; Nakamura et al., 2016). These studies have yielded broad overviews of epigenetic status and transcriptional activity in early embryonic development.

To date, three reports provide single-cell RNA-seq data from human embryos to the blastocyst stage, entailing a total of 1683 individual transcriptomes (Yan et al., 2013 (*n*=124); Blakeley et al., 2015 (*n*=30); Petropoulos et al., 2016 (*n*=1529)). The studies differ in scope of developmental timespan, sample collection and processing, sequencing protocol and cell-type classification. Such differences present substantial analysis challenges that impair direct comparison of embryonic lineages between datasets (Vallot et al., 2017). The accurate and consistent distinction of EPI and PrE cells has proven particularly difficult and consequently impedes the analyses of fate specification events.

Here we undertake a comprehensive analysis of human embryo single-cell transcriptome datasets, characterising the features of each and resolving ambiguities in cell-type assignments. We build a unified transcription map comprising representative samples of defined embryonic lineages. The associated gene expression signatures recapitulate known lineage marker protein localisation in embryos assayed by immunofluorescence, and enable the discovery of specific transcriptional events and regulatory networks over developmental time. This study provides detailed insight into the emergence of pluripotency in the human embryo and an analytical framework in which to assess the developmental state of self-renewing pluripotent cell lines cultured ex vivo.

## Results

### TE overrepresentation in single-cell embryo data

We embarked on a systematic analysis of single-cell transcriptome data from three human embryo profiling studies that extend to late blastocyst (Yan et al., 2013; Blakeley et al., 2015; Petropoulos et al., 2016). When examining the most extensive dataset produced to date (Petropoulos et al., 2016), global analysis of EPI and PrE cells as classified in that study showed substantial overlap between the two populations (Fig. 1A, S1A). Alternative dimensionality reduction algorithms, such as t-SNE (Fig. S1B) and diffusion map visualisation methods (Fig. S1C), generated consistent sample clusters. Moreover, lack of separation between EPI and PrE cells could not be attributed to potential differences in samples collected at E6 and E7 (Fig. S1D). A degree of transcriptional identity may be expected as it has been noted that EPI and PrE share broadly similar expression profiles at early blastocyst stages in the mouse embryo, and are most effectively distinguished by a subset of marker genes (Boroviak et al., 2015). However, significant contribution from genes enriched in TE was observed in the first dimension of principal component analysis (PCA) (Fig. 1B). Thus when the entire dataset is considered as an ensemble, the presence of TE cells impedes accurate resolution of EPI and PrE and subsequent characterisation of those lineages.

**Fig. 1.**
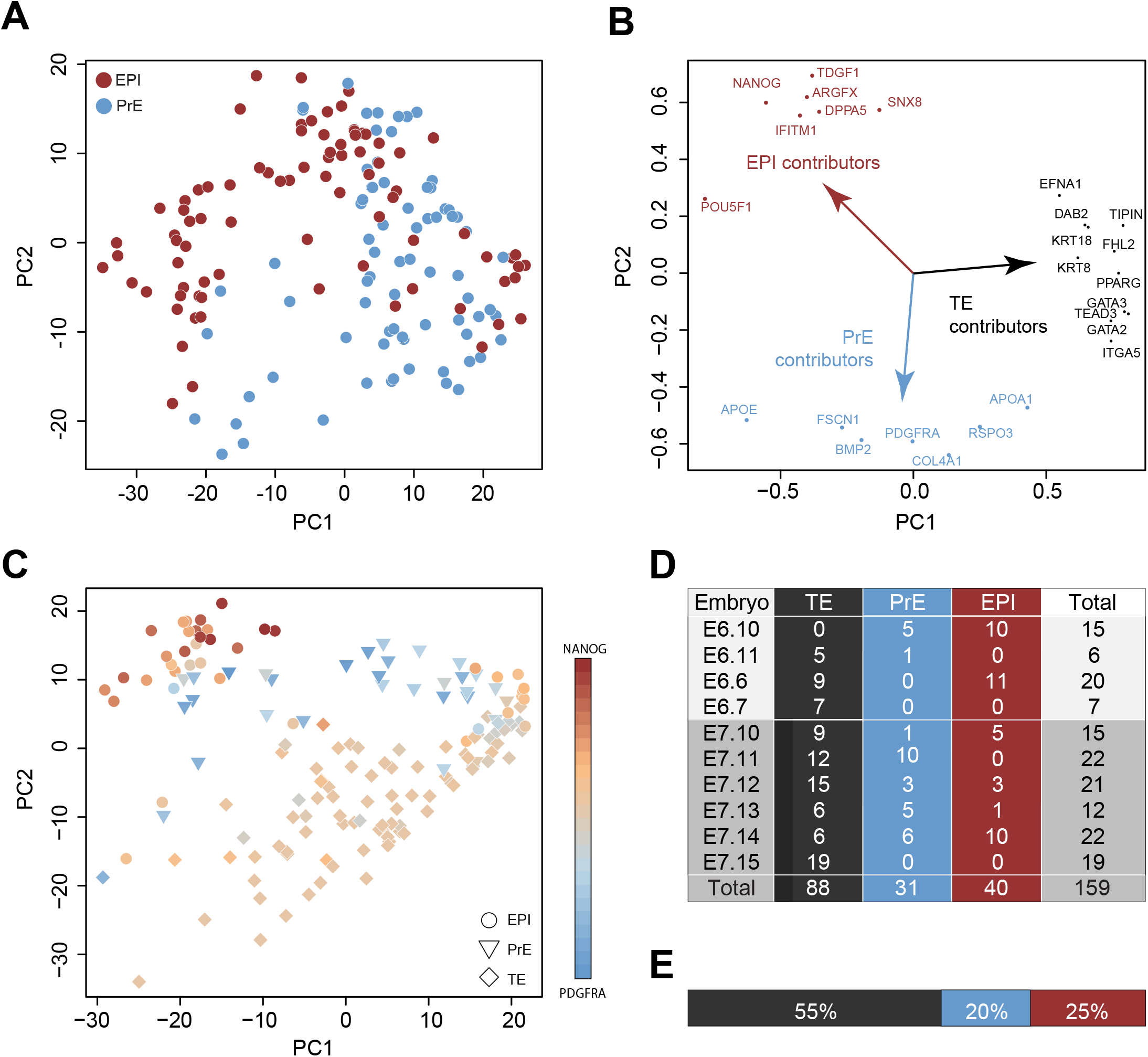
Embryo lineage classification. (A) PCA of E6 and E7 samples based on the most variable genes (*n*=1294, log_2_ FPKM > 2, log CV > 0.5), coloured according to cell type classification by Petropoulos et al. (2016). (B) Genes contributing to the first and second principal components. (C) PCA of E6 and E7 immunosurgery samples from Petropoulos et al. based on variable genes (*n*=1131, log_2_ FPKM > 2, log CV^2^ > 0.5). Colours are scaled to the ratio of *NANOG* (EPI) to *PDGRA* (PrE) expression. (D) Lineage assignments of E6 and E7 immunosurgery samples according to Petropoulos et al. (E) Relative percentages of EPI, PrE and TE cells from embryos processed by immunosurgery as reported by Petropoulos et al.

A subset of samples from Petropoulos et al. was obtained from embryos treated by immunosurgery, which canonically entails ablation of the TE by complement-mediated cell lysis and mechanical isolation of intact ICM (Solter and Knowles, 1975). To determine the properties of EPI and PrE lineages in a dataset presumed to be devoid of TE cells, we examined those samples captured via immunosurgery from late blastocysts at E6 and E7. At this stage, EPI and PrE are largely discerned by marker analysis (Roode et al., 2012; Niakan and Eggan, 2013). However, PCA based on the most variable genes did not yield distinct EPI and PrE populations (Fig. 1C). Plotting the ratio of *NANOG* (EPI) versus *PDGFRA* (PrE) expression revealed an EPI population comingled with a minority of PrE cells, but the largest proportion displayed intermediate levels of *NANOG* and *PDGFRA* (Fig. 1C). The predominant genes contributing to the separation of samples were TE-associated, including *GATA2, GATA3, KRT8, DAB2* and *TEAD3* (Fig. S1E). Indeed, many of the cells concerned were classified as TE in the primary report (Petropoulos et al., 2016). Samples were derived from four E6 and six E7 embryos (Fig. 1D) and more than half were annotated to belong to the TE lineage (Fig. 1E). This is highly unexpected and suggests incomplete immunolysis and ICM recovery in the original study.

### Lineage markers defining human EPI, PrE and TE

We sought to compile a robust dataset of representative EPI and PrE transcriptomes from available single-cell profiling data. Ideally, this dataset should contain samples from each of the three published studies (Yan et al., 2013; Blakeley et al., 2015; Petropoulos et al., 2016) and recapitulate known lineage marker localisation (Kuijk et al., 2012; Roode et al., 2012; Niakan and Eggan, 2013; Blakeley et al., 2015; Deglincerti et al., 2016; Guo et al., 2016). We assembled a set of 12 high-confidence marker genes described in the literature, four associated with each of the three blastocyst lineages (Fig. 2A). We evaluated the discriminatory power of these genes on cells profiled in the Yan and Blakeley studies (Fig. 2B,C). We found that clear separation between EPI, PrE and TE could be attained for nearly all samples. This result indicates that post hoc identification of early human embryo cells based on this minimal set of lineage markers is compatible with the cell-type classification proposed in Blakeley et al. (Fig. S2A, Table S1), and further confirms those assignments as consistent with published immunofluorescence data.

**Fig. 2.**
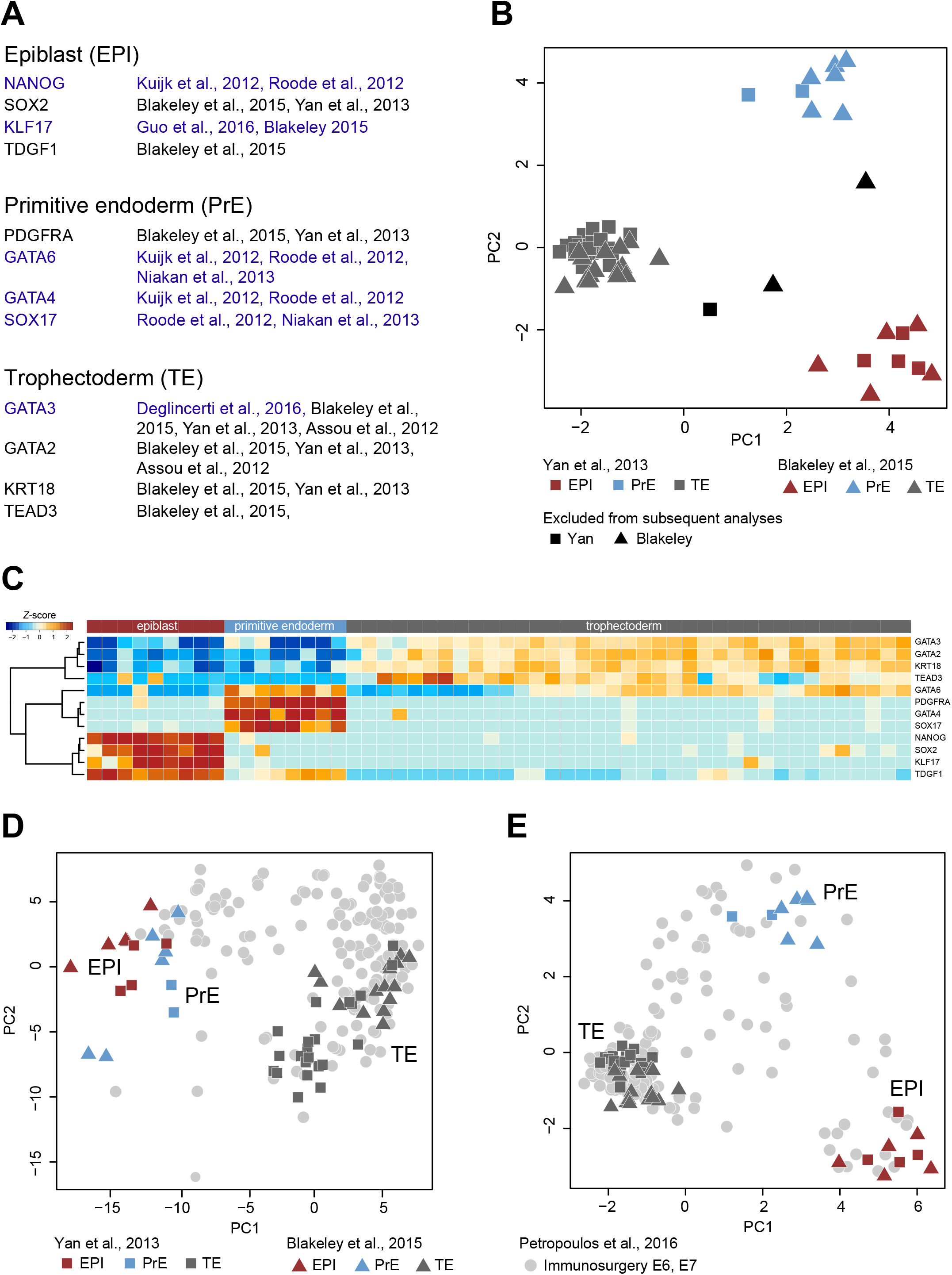
Lineage segregation based on marker genes. (A) Panel of 12 high-confidence markers for EPI, PrE and TE. Publications with immunofluorescence data showing protein expression in the human blastocyst are highlighted in blue. A subset of TE cells express CDX2 in human (Chen et al., 2009; Niakan et al., 2012). (B) PCA of embryo cells profiled in the Yan and Blakeley studies based on lineage markers. (C) Hierarchical clustering of Yan and Blakeley datasets. (D) PCA of EPI, PrE and TE with E6 and E7 Petropoulos immunosurgery samples (grey) based on common variable genes (*n*=188, log_2_ FPKM > 2, log CV^2^ > 0.5). (E) PCA as in D based on lineage markers.

Upon inclusion of E6–E7 EPI and PrE samples as annotated in Petropoulos et al., we observed substantial overlap between EPI, PrE and TE cells from all three studies, with limited dispersion of Petropoulos samples that precluded resolution of distinct lineages (Fig. S2B). PCA of a combined dataset comprising Blakeley and Yan cells with the Petropoulos immunosurgery subset revealed substantial apparent technical bias between the Petropoulos study and others, largely captured by PC1 (Fig. S2C,D). To mitigate these differences and allow meaningful comparisons between studies, we identified the most variable genes in each dataset (Fig. S2E). Intersecting these yielded 188 genes common to all (Fig. S2F). PCA confined to this gene set showed a substantial reduction in technical bias (Fig. 2D). This analysis further confirmed that the majority of immunosurgery-derived samples from Petropoulos et al. likely originate from TE. PCA based on lineage markers produced similar results (Fig. 2E).

### Representative transcriptomes of mature EPI and PrE lineages

To date the Petropoulos study embodies the most extensive human embryo single-cell transcriptome resource, comprising 858 late blastocyst samples among 1529 total sequenced. To discriminate presumptive EPI and PrE from TE cells in this dataset, we examined POU5F1 (OCT4) expression at the late blastocyst stage. Immunofluorescence assays have tracked POU5F1 expression from E3 (8-cell) to E7 (late blastocyst), where localisation is nonspecific in all cells of the compacted morula and spans most cells of the early blastocyst (Fig. 3A, (Niakan and Eggan, 2013)). From the mid-blastocyst stage, POU5F1 is gradually restricted to the ICM. Notably, POU5F1 is expressed in the ICM throughout preimplantation development (Niakan and Eggan, 2013).

**Fig. 3.**
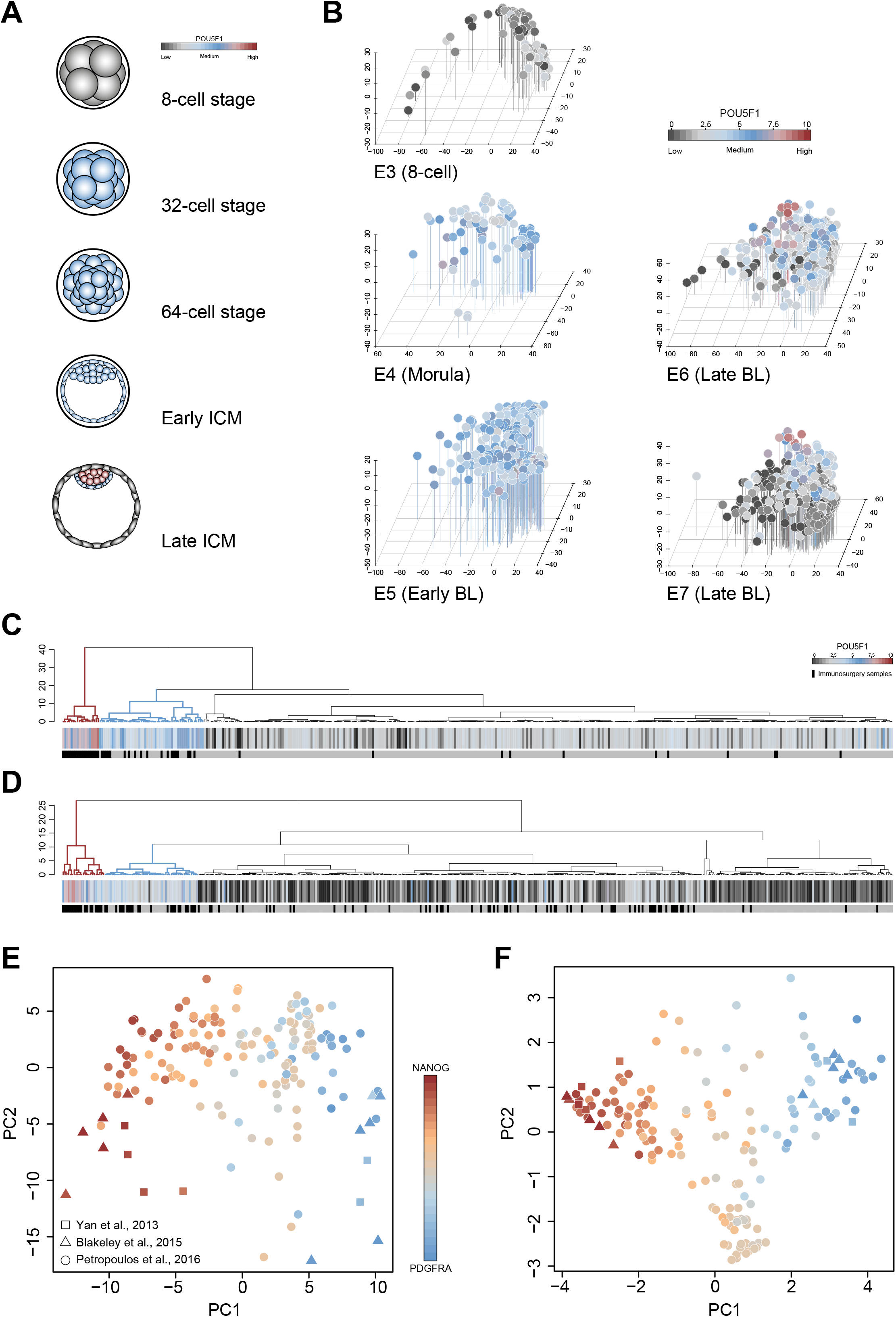
Identification of late ICM cells. (A) Schematic of preimplantation development from the 8-cell stage according to published immunofluorescence assays (Niakan and Eggan, 2013). Colours depict POU5F1 localisation typical of each stage. (B) PCA of Petropoulos samples based on variable genes (log_2_ FPKM > 2, log CV > 0) according to embryonic day. Colours represent log_2_ *POU5F1* expression. (C–D) Dendrograms of E6 (C) and E7 (D) Petropoulos samples derived from the third principal component in B, indicating *POU5F1* expression. POU5F1-high and -medium clusters are highlighted in red and blue, respectively. Bars indicate individual cells with those recovered by immunosurgery indicated in black. (E) PCA of Petropoulos samples based on common variable genes (*n*=170, log_2_ FPKM > 2, log CV^2^ > 0.5) with high and medium *POU5F1* levels as selected in C and D, together with EPI and PrE cells from Yan and Blakeley datasets. (F) PCA based on marker genes as described in E.

To determine if similar dynamics were evident at the transcript level, we examined *POU5F1* expression across developmental stages (Fig. 3B, S3A,B). *POU5F1* mRNA levels closely resembled the pattern of protein abundance observed throughout the embryo as established by immunostaining. At mid to late blastocyst stages E6 and E7 *POU5F1* downregulation was seen in the majority of cells, apart from a small population representing the late ICM (Fig. 3B, S3A,B). To extract this set we performed hierarchical clustering based on the third principal component to distinguish POU5F1-high from *POU5F1-low* cells (Fig. 3C,D). This facilitated isolation of clusters of E6 (Fig. 3C) and E7 (Fig. 3D) cells displaying medium and high *POU5F1* expression. PCA based on highly variable genes (Fig. 3E) and lineage markers (Fig. 3F) revealed close correspondence of E6 and E7 POU5F1-high cells to EPI and PrE samples from the Blakeley and Yan datasets. EPI and PrE cells segregated along PC1, suggesting that separation of these groups is largely attributed to biological differences between the two lineages. Exclusive selection of POU5F1-high clusters (Fig. S3C,D) produced similar results, although we noted a slight underrepresentation of PrE cells (Fig. S3E,F). This is consistent with expression patterns observed in the late human blastocyst, where POU5F1 appears to be higher in EPI than in PrE (Roode et al., 2012). We conclude that *POU5F1* mRNA levels recapitulate previously observed protein localisation. Thus identification of POU5F1-high and -medium expressing sample groups facilitates a posteriori separation of ICM from TE cells.

Hierarchical clustering of POU5F1-high and -medium cells at E6 and E7, together with EPI and PrE samples from the Yan and Blakeley datasets, produced three distinct populations (Fig. 4A). *NANOG:PDGFRA* expression ratios in these cells are indicative of EPI, PrE and an intermediate group. We derived transcriptomes representing mature EPI and PrE lineages as well as intermediate cells from the associated clusters (Fig. 4A), comprising data from all studies considered. EPI and PrE samples formed discrete groups by genome-wide expression (Fig. S4A) and marker-based PCA (Fig. S4B). According to this classification all EPI and PrE cells exhibited robust expression of high-confidence lineage markers, indicating mature EPI and PrE identity (Fig. 4B). Intermediates were characterised by mid-range levels of *GATA6* and heterogeneous expression of both EPI and PrE markers (Fig. 4B). We infer that these intermediates may represent ICM cells in the process of transition to either lineage. Nonetheless, a substantial number of ICM cells have advanced to mature EPI and PrE identities and readily resolve into distinct populations, a finding that becomes apparent when our selection process is applied (Fig. S4C,D). Thus, subpopulations of EPI and PrE cells have acquired distinct fates by the late blastocyst stage while other ICM cells appear to be transitory.

**Fig. 4.**
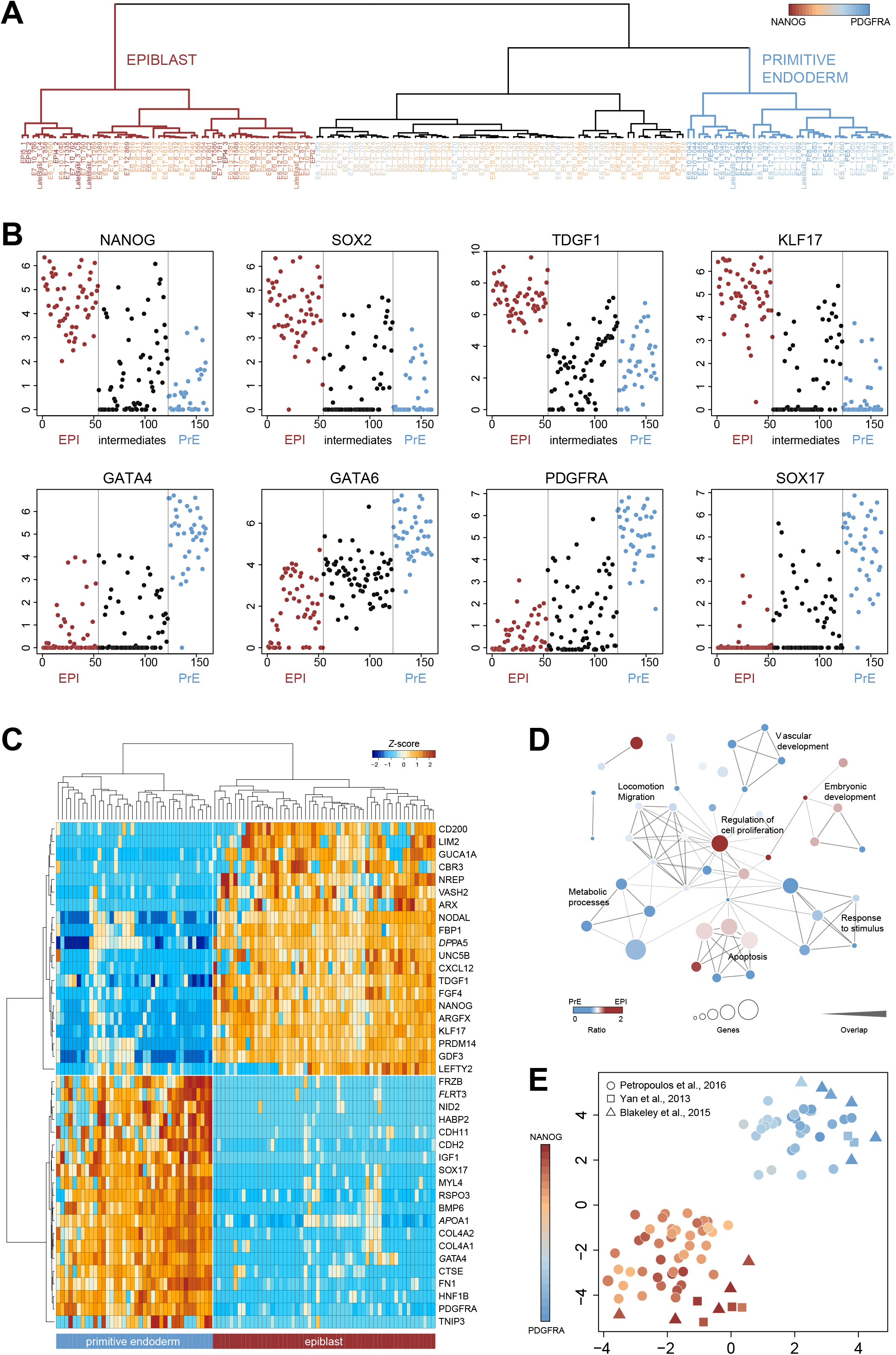
Selection and characterisation of EPI and PrE cells. (A) Cluster dendrogram of *POU5F1-high* and -medium expressing late ICM cells, selected based on the first and second principal components from the analysis in Fig. 3F. Sample colours are scaled to the ratio of *NANOG* to *PDGFRA* expression. (B) Single-cell dot plots showing log_2_ FPKM values of the genes indicated. Cells are ordered along the *x*-axis to correspond to the dendrogram in A. (C) Two-way clustering of the top 20 genes up- and down-regulated between EPI and PrE samples. (D) Network of biological processes enriched for genes modulated between EPI and PrE. Nodes represent processes; edge weight reflects the degree of intersection between gene lists. Node size is proportional to the number of contributing genes and colours reflect the ratio between those up- and down-regulated. (E) t-SNE plot for EPI and PrE cells. Sample colours are scaled to the ratio of *NANOG* to *PDGFRA* expression.

We identified additional lineage-associated genes consistently expressed in the corresponding cell types. (Fig. S4E, F). Primate-specific PrE marker *RSPO3* (Boroviak et al., 2015) was upregulated in reclassified PrE cells, while EPI cells featured pluripotency markers *PRDM14, TFCP2L1* and *ZFP42*. Mouse-specific factors of the pluripotency network, including *KLF2, NR0B1* and *FBXO15* (Blakeley et al., 2015; Boroviak et al., 2015) were absent. Differentially expressed genes were then identified between EPI and PrE (Fig. 4C, Table S2). Consistent with previous observations, we found primate-specific EPI expression of *ARGFX, NODAL* and *LEFTY2,* whereas components of BMP signalling were evident in PrE cells (Blakeley et al., 2015; Boroviak et al., 2015; Petropoulos et al., 2016). Gene Ontology (GO) analysis of transcripts modulated between EPI and PrE populations showed enrichment for apoptosis, cell proliferation and embryo development in EPI cells. PrE-related processes included cell migration, metabolism and vascular development (Fig. 4D). Finally, we assessed the purity of EPI and PrE samples with t-SNE based on lineage markers. We obtained tight separation between the two groups (Fig. 4E), indicating they comprise distinct and representative populations of mature EPI and PrE cells.

### Pseudotime staging and non-human primate-derived marker expression define the early ICM

Cells from early blastocyst (E5) embryos were profiled only in the Petropoulos study. PCA suggested the presence of a heterogeneous mixture of cells (Fig. 5A). Combining developmental pseudotime (Fig. 5A) with hierarchical clustering (Fig. S5A) allowed us to resolve an early cluster comprising 43 of 336 E5 cells (Fig. 5A, early ICM). In agreement with Petropoulos et al., we could not detect an EPI, PrE or TE signature in this population (Fig. 5B). Instead we observed intermediate levels of both *SOX2* and *GATA3* (Fig. S5B), indicating a subset of cells expressing a mixture of ICM and TE lineage markers. Analysis of genes contributing to the first and second principal components revealed an early TE population displaying high levels of *GATA2, GATA3* and *DAB2* (Fig. S5C). ICM cells expressing *IFITM1, GDF3* and *ARGFX* formed a separate group, consistent with the cell types assigned by Petropoulos et al. (Fig. 5B, S5C).

**Fig. 5.**
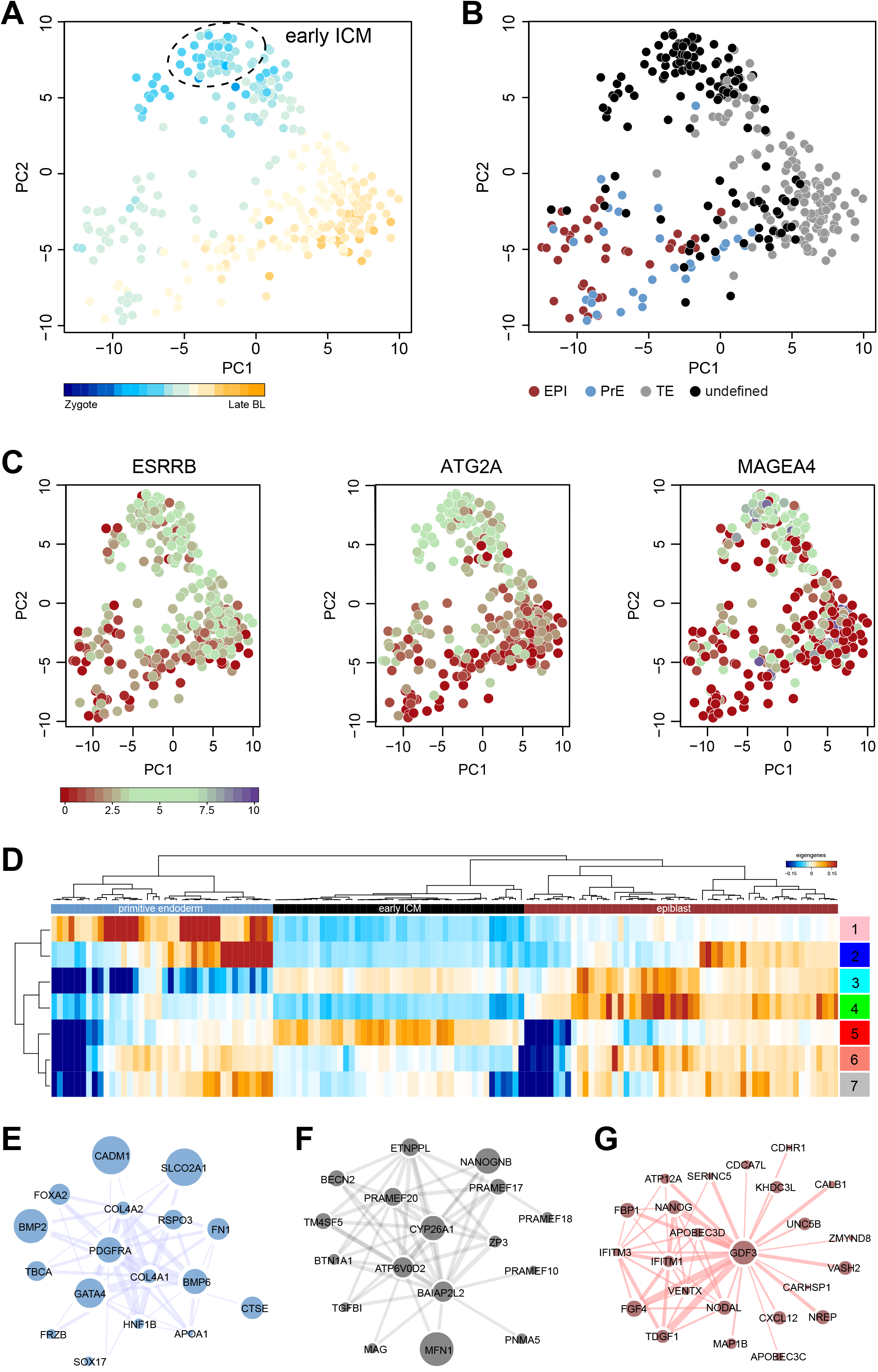
Identification of early ICM and WGNCA analysis of lineage segregation. (A–C) PCA based on highly variable genes (*n*=203, log_2_ FPKM > 2, log CV^2^ > 1) for Petropoulos E5 cells. Samples are coloured by (A) pseudotime, (B) lineage classification according to Petropoulos et al. and (C) absolute expression in log_2_ FPKM for early ICM markers in the common marmoset and rhesus macaque. (D) Two-way clustering of eigengene values as computed by WGCNA. The first three major branches corresponded to PrE (Module 1), early ICM (Module 5) and EPI (Module 4). (E–G) Networks of highly co-regulated genes in PrE (E), early ICM (F) and (G) EPI lineages. Node sizes are proportional to absolute fold change in expression between EPI and PrE or to absolute expression for early ICM. Edge thickness reflects the number of co-regulated genes between adjacent nodes.

To date defined markers of the early human ICM have not been proposed. In studies of non-human primates, *ESRRB* has recently been identified as an early ICM-specific gene in cynomolgus macaque (Nakamura et al., 2016) and common marmoset (Boroviak et al., in preparation). *ESRRB* is robustly expressed in the putative human early ICM population and down-regulated in EPI, together with other early ICM-specific genes *ATG2A* and *MAGEA4* (Fig. 5C). Inclusion of “early E5” cells from the Petropoulos study, collected several hours prior to those designated E5 by IVF staging (Fig. S5D), resulted in substantial overlap with the early ICM population. This result independently confirmed cell-type assignments inferred by pseudotime analysis.

To further assess whether early ICM, EPI and PrE cells can be discerned as separate populations, we performed weighted gene network cluster analysis (WGCNA, Zhang and Horvath, 2005) based on highly variable genes (Fig. S5E). Using this approach we extracted gene modules defined by co expression combined with unsupervised clustering. We identified 14 initial modules, which could be reduced to seven based on similarity metrics (Fig. S5F). Importantly, eigengene clusters based on Pearson correlation independently captured our previously refined cell populations. We used the 50 most highly connected genes to define co-expression networks for PrE (Fig. 5E), ICM (Fig. 5F) and EPI lineages (Fig. 5G) (Table S3). *PDGFRA, BMP6, RSPO3, COL4A1* and *GATA4* were the major hub genes for the PrE network, whereas *GDF3,* together with *NANOG, IFITM1, IFITM3, TDGF1* were highly co-expressed in the EPI module. Predominant hub genes in the early ICM network consisted of *MFN1, CYP26A1, NANOGNB, PRAMEF17* and *PRAMEF20.* These results demonstrate that early ICM, EPI and PrE can be resolved as distinct transcriptional states.

### A unified transcription map of human preimplantation development

We then integrated samples from earlier developmental stages spanning zygote to compacted morula (Yan et al., 2013), analysing these in tandem with the refined classifications of early ICM, EPI and PrE established above (Table S4, S5). PCA produced stage-specific clusters (Fig. 6A), verifying that the cell populations established here reflect distinct embryonic lineages. We then generated self-organising feature maps (Kohonen, 1982) from this combined dataset to extract the most prominent stage-specific transcription factors, chromatin modifiers and biological processes operative in human preimplantation development (Fig. 6B, Table S6). Genes were ranked by expression level and *Z*-score which in turn captured specific embryo stages as an emergent property. Embryonic genome activation at the 8-cell stage was characterised by RNA metabolic processes and expression of *LEUTX,* a homeobox gene recently associated with this event and without an orthologue in mouse (Maeso et al., 2016). Expression of *KLF17* also peaked at the 8-cell stage. We found *ZNF296*, encoding a pluripotency-associated protein reported to interact with the reprogramming factor Klf4 in mouse (Fischedick et al., 2012; Fujii et al., 2013; Matsuura et al., 2017), specifically expressed in compacted morulae. Five of the top ten genes upregulated at the late ICM stage are mutually exclusive to EPI (*LEFTY2, IFITM1* and *LEFTY2*) or PrE (*APOA1* and *RSPO3*) lineages.

**Fig. 6.**
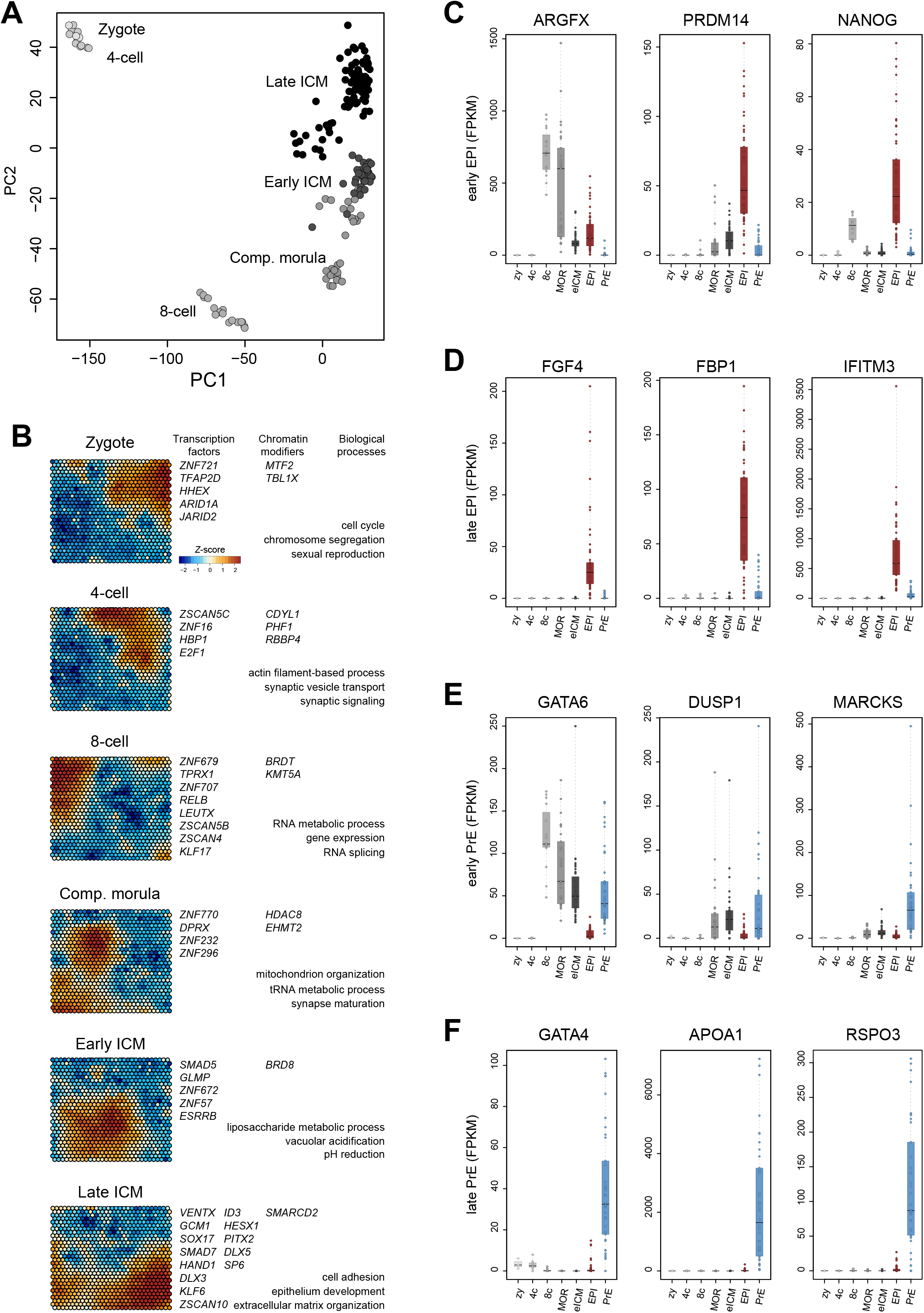
WGCNA and global analysis of human preimplantation development. (A) PCA of zygote, 4-cell, 8-cell and compacted morula samples from the Yan dataset combined with our selection of early ICM cells from Petropoulos et al. and the refined subset of EPI and PrE from all three studies. (B) Self-organising maps of selected transcription factors, chromatin modifiers and significant biological processes across developmental stages. (C–F) Expression in FPKM of selected markers for (C) early EPI, (D) late EPI, (E) early PrE and (F) late PrE.

We sought to identify progressive drivers of EPI and PrE specification in human (Fig. 6C–F). Transcriptional regulators *ARGFX, PRDM14, SOX2, NANOG* and *DPPA2* were expressed from the point of embryonic genome activation and maintained in the EPI, but downregulated in PrE (Fig. 6C, S6A). Strikingly, EPI-specific genes, and specifically those upregulated in mature EPI cells, included several agonists of FGF, Activin A/Nodal and WNT signalling, such as *FGF4, GDF3, NODAL, LEFTY2, TDGF1* and *WNT3* (Fig. 6D, S6B). Notable early PrE-associated genes were *GATA6, LAMA1, HNF4A, DUSP1* and *MARCKS* (Fig. 6E, S6C). In contrast, *RSPO3, GATA4, APOA1, SOX17, FOXA2, APOA2, PDGFRA* and *BMP2* were upregulated de novo in the mature PrE (Fig. 6F, S6D). This molecular classification of early and late EPI and PrE lineages (Table S7) is thus a means to stage human preimplantation development and provides a basis for functional interrogation.

### Characteristics of human pluripotent stem cell lines

Pluripotent stem cells (PSC) can be derived from explant cultures of human blastocyst ICM (Thomson et al., 1998; O’Leary et al., 2012) and propagated ex vivo in the presence of FGF and Activin or TGFß (James et al., 2005; Vallier et al., 2005; Xu et al., 2005). Having established a defined sequence of early human embryo progression, we sought to relate the transcriptional status of PSC cultured in vitro to embryonic lineages. We compiled an extensive dataset of human PSC lines cultivated in standard conditions (see Methods), several modalities that support the transition from conventional PSC to a naïve state with transcriptomic, epigenetic and metabolic features similar to canonical mouse embryonic stem cells (Takashima et al., 2014; Theunissen et al., 2014; Guo et al., 2016; Guo et al., 2017), alternative methods where the presumption of a naïve identity has been advanced but not corroborated (Gafni et al., 2013; Ware et al., 2014), and a recent report describing cells proposed to possess extended developmental potential (Yang et al., 2017).

PCA (Fig. 7A), hierarchical clustering (Fig. S7A) and t-SNE (Fig. S7B) consistently partitioned samples into two broad groups, unambiguously distinguishing naïve from conventional and alternative cultures. The naïve cluster exclusively comprised cells reported in four studies: conversion of conventional human PSC to a naïve state using 5i/L/A (MEKi, GSK3βi, BRAFi, SRCi, ROCKi, LIF, Activin A) (Theunissen et al., 2014) and three studies employing t2iLGö (MEKi, GSK3i, LIF, PKCi) in transgene-mediated resetting (Takashima et al., 2014), derivation of de novo cell lines directly from dissociated ICM cells (Guo et al. 2016), and chemical resetting (cR) (Guo et al. 2017). Included in the last dataset were cells cultivated in feeder-free conditions on a laminin attachment substrate. These clustered with other samples in the naïve group (Fig. 7A, S7B).

**Fig. 7.**
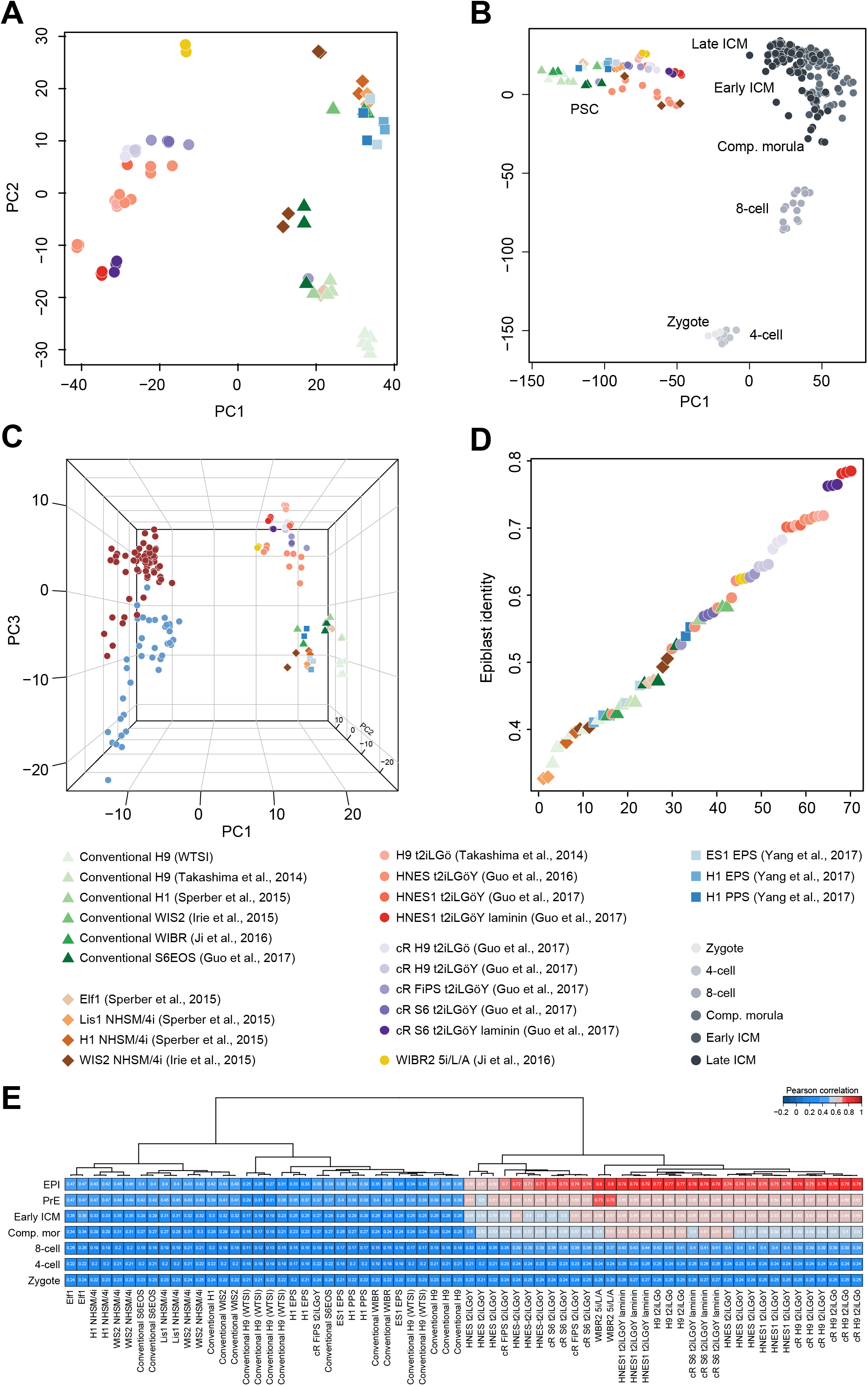
Comparison of human pluripotent cell lines to embryonic stages. (A) PCA based on highly variable genes (*n*=1760, log_2_ FPKM > 2, log CV^2^ > 0.5). (B) PCA based on global gene expression for all preimplantation stages and PSC cultures. (C) PCA of late blastocyst stages (red=EPI, blue=PrE) and PSC cultures. (D) Fraction of identity of PSC cultures to EPI cells. Similarity between cultured PSC and all preimplantation embryo stages was computed by quadratic programming; plotted is the fraction identity of PSC to EPI, with samples sorted accordingly. (E) Clustering of Pearson correlation of genes differentially expressed between naïve and conventional PSC to embryo stages and lineages (*n*=2860, adjusted *p* < 0.001, absolute log_2_ fold change > 1.5).

PSC lines maintained in modified NHSM/4i and serum replacement factors (MEKi, GSK3i, LIF, FGF2, TGFβ1, p38i, JNKi, ROCKi, KSR) (Irie et al., 2015; Sperber et al., 2015), or generated by transient exposure to HDAC inhibitors followed by propagation in alternative media (MEKi, GSK3i, LIF, FGF2, IGF1, ROCKi, KSR) (Sperber et al., 2015) were interspersed with those cultured in conventional conditions by global dimensionality reduction methods (Fig. 7A, S7A,B). These results are consistent with previous studies in which the relationship of these cells to standard PSC was assessed (Huang et al., 2014; Nakamura et al., 2016; Pastor et al., 2016). Statistical testing identified “Preimplantation Embryo” as the topmost ranked functional pathway based on genes enriched in the naïve sample cluster, whereas “Ectoderm Differentiation” was most associated with conventional PSC. This is in agreement with reports suggesting that naïve stem cells represent an early stage of embryonic development (Huang et al., 2014; Theunissen et al., 2016) whereas conventional primate PSC more closely resemble the postimplantation epiblast (Nakamura et al., 2016).

Using the composite single-cell embryo dataset as a reference, we then explored the relationship between human PSC cultures and embryonic lineages over developmental time. Initial clustering by PCA indicated sample processing methods to be the main contributor to PC1, likely arising from technical factors inherent to single-cell and bulk RNA-seq protocols (Fig. 7B). PC2 resolved developmental timing, and aligned in vitro cultured PSC with ICM stages. This result was recapitulated by alternative global dimensionality reduction methods (Fig. S7C). To increase the resolution of cluster separation, we replotted PSC datasets with compacted morula, early ICM and late ICM embryo samples (Fig. S7D). PSC were invariably observed in closest proximity to late ICM. Confining the analysis to late ICM and in vitro samples alone resolved distinct groups of EPI and PrE embryo cells, and separated naïve and conventional PSC cultures (Fig. 7C). Notably, naïve PSC aligned with EPI cells along the third dimension.

As an alternative approach, we employed quadratic programming to compare cell expression profiles between PSC and embryonic tissues based on the entire transcriptome (Gong and Szustakowski, 2013). This allowed us to compute fractional identity between PSC cultures and preimplantation embryo lineages (Fig. 7D). Naïve PSC showed the greatest shared identity with EPI cells, over 0.75 for laminin cultures, in contrast to conventional PSC and other alternatives which consistently displayed lower correspondence (<0.54). We also performed correlation analysis based on genes dynamically expressed over developmental time (Fig. S7E). We observed a gradual increase in correlation with embryonic progression, most prominently with the EPI subset.

Finally, we compared in vitro cultured cells to the embryo based on genes differentially expressed between naïve and conventional PSC sample clusters (Table S9). The two groups were clearly separated, where naïve samples most closely resembled EPI cells (Fig. 7E; 0.66–0.8 for naïve vs 0.19–0.44 for conventional PSC). Interestingly, reset cells (Takashima et al., 2014) were also correlated to varying degrees with PrE (0.61–0.73), early ICM (0.54–0.67) and compacted morulae (0.5–0.64). Cells adapted to 5i/L/A (Ji et al., 2016) exhibited the highest correlation (0.8) with both EPI and PrE. Cultures of embryo-derived (Guo et al., 2016) and chemically reset (Guo et al., 2017) cells in t2iLGö were similarly correlated with EPI as 5i/L/A cells, but to a lesser extent with PrE. These findings indicate that human PSC maintained in t2iLGö represent cogent transcriptional counterparts to the naïve epiblast lineage of the human preimplantation embryo and suggest that 5i/L/A cells may represent a mixed EPI/PrE identity.

## Discussion

Here we present an analysis and classification of single-cell gene expression data from human embryos, and define accurate transcriptional prototypes of EPI and PrE lineages emergent in the late blastocyst. We integrated data from disparate profiling studies and determined representative cell populations that distinguish nascent and differentiating tissues. The resulting dataset is consistently partitioned by developmental stage and cell type using multiple dimensionality reduction methods, and the constituent cell populations faithfully recapitulate expression and localisation patterns of established embryonic lineage markers.

WGCNA independently clustered samples into groups consisting of early ICM, EPI and PrE. This partitioning allows further exploration of specific gene sets, including those encoding extracellular matrix proteins and signalling pathway components, transcriptionally active during progression from unspecified ICM to mature EPI and PrE lineages. EPI cells lack expression of several genes active in mouse EPI and ESC, such as *ESRRB, FBXO15, NR0B1* and *KLF2,* consistent with previous reports in human (Blakeley et al., 2015; Petropoulos et al., 2016) and non-human primates (Boroviak et al., 2015; Nakamura et al., 2016). The human naïve pluripotency network includes transcription factors *MYBL2, ARGFX, SOX4, PRDM14, KLF4, TFCP2L1, GDF3* and *KLF17.* We identify markers of early lineage EPI and PrE specification, many of which are expressed from embryonic genome activation at the 8-cell stage, such as *KLF17* and *ARGFX.* The temporal sequence of human PrE transcription factor acquisition was surprisingly well conserved to the rodent paradigm (Plusa et al., 2008; Rossant and Tam, 2009; Artus et al., 2011; Schrode et al., 2014). *GATA6* preceded *SOX17,* followed by *GATA4* and *FOXA2* in the ICM; however we did not detect PrE-specific expression of *SOX7.*

Immunofluorescence studies of the human blastocyst have shown that EPI and PrE segregation is manifest prior to implantation, between E6 and E7 (Roode et al., 2012; Niakan and Eggan, 2013; Deglincerti et al., 2016). Similar observations have been made in non-human primates. The early marmoset ICM co-expresses GATA6 and NANOG, before the point where EPI and PrE lineages diverge (Boroviak et al., 2015). In cynomolgus monkey, the early ICM comprises a distinct population that subsequently undergoes EPI and PrE segregation at the late blastocyst stage, similar to mouse (Nakamura et al., 2016; Ohnishi et al., 2014). Consistent with these findings, our analyses support a discrete developmental state embodied by the early human ICM, different in composition from mature EPI or PrE.

Petropoulos et al. instead proposed concurrent establishment of EPI, PrE and TE lineages during blastocyst formation at E5. An overlap between acquisition of ICM versus TE identity and EPI and PrE specification is not excluded by our analysis, but in that event, divergence of EPI and PrE would commence in a subset of ICM cells at E5 and progress incrementally through E6. Segregation of EPI and PrE appears ongoing for at least one cell division cycle in mouse (Saiz et al., 2016). Consistent with the hypothesis of Petropoulos et al., we identified a fraction of E5 cells that express lineage-specific markers. However, it is conversely plausible that these cells were obtained from more advanced embryos. Staging of human embryos is not as precise as the rodent model, and it could be that some human IVF-derived blastocysts are more or less developmentally mature than embryonic day would imply.

The revised lineage assignments presented here clearly demarcate late ICM populations into EPI, PrE and putative transitional intermediates observed between EPI and PrE in the late blastocyst. The fate of these intermediates remains an open question. Pathway analysis of prototypical EPI cells revealed significant enrichment for apoptosis, which may facilitate selective elimination of unspecified cells. Alternatively, these cells may persist for some time before commitment to a particular lineage.

Human and non-human primate PSC propagated in vitro by conventional culture methods differ substantially from pluripotent cells resident in the preimplantation embryo with respect to transcriptome (Yan et al., 2013; Nakamura et al., 2016) and methylome (Guo et al., 2014). Conventional self-renewing PSC lines share distinguishing features with EpiSC derived from mouse postimplantation epiblast (Brons et al., 2007; Tesar et al., 2007), which has led to the proposition that these cultures may have progressed in vitro towards a later stage of development (Nichols and Smith, 2009; Davidson et al., 2015). A genuine primate analogue to rodent ESC has been sought in recent years. That goal has remained elusive, and it has been unclear whether suboptimal culture conditions and/or species differences in pluripotency networks impeded the capture of human naïve cells with properties similar to authentic ESC.

The degree to which various in vitro methods promote the conversion of human PSC to a naïve state has been assessed in terms of global transcription, induction or suppression of pluripotency-associated and lineage marker genes, activation of retroviral element families, genome-wide DNA methylation, X chromosome activation, and other properties (Huang et al., 2014; Pastor et al., 2016; Nakamura et al., 2016; Theunissen et al., 2016, Liu et al., 2017). The sequence of human preimplantation development we have derived allows comprehensive transcriptome comparison of embryo stages to PSC propagated in various culture systems. In agreement with earlier reports, we found that independent samples of PSC maintained in NSHM/4i (Irie et al., 2015; Sperber et al., 2015) did not appreciably diverge from conventional cultures in transcriptional state. Extended pluripotent stem (EPS) cells (Yang et al., 2017) were correlated with conventional PSC on a global level and not with any preimplantation embryo stage, suggesting that any altered potential may be an in vitro adaptation conferred by a relatively small-scale regulatory perturbation.

In contrast to the above, cells converted to a naïve state via exogenous transgene expression or chemical resetting and propagated in t2iLGö (Takashima et al., 2014; Guo et al., 2017), de novo embryo-derived naïve PSC lines established in similar conditions (Guo et al., 2016), and PSC maintained in 5i/L/A (Theunissen et al., 2014) retained strong correspondence to cells of the preimplantation embryo. These naïve pluripotent cultures were also variably correlated to PrE, early ICM and compacted morula. Mouse PrE and EPI display global transcriptional similarity despite differential expression of lineage specifiers (Boroviak et al., 2015). The apparent concordance between naïve PSCs and PrE is probably rooted in the intrinsic global similarity between EPI and PrE.

However, the strongest degree of similarity was observed between naïve cultures and EPI. The correspondence was closest for 5i/L/A cultures, and for feeder-free HNES cells and chemically reset PSC in t2iLGö. We therefore conclude these culture regimes capture self-renewing pluripotent cell populations characterised by transcriptomes that approximate to the in vivo preimplantation epiblast.

## Materials and methods

### RNA-seq data processing

Sequencing data were obtained from the European Nucleotide Archive (Toribio et al., 2017) from single-cell human embryo profiling studies (accessions SRP011546 (Yan et al., 2013), SRP055810 (Blakeley et al., 2015), ERP012552 (Petropoulos et al., 2016)), H1 (SRP014320 (Djebali et al., 2012)) and H9 (ERP007180 (Wellcome Trust Sanger Institute)) PSC lines cultured in standard conditions, H9 cells in conventional conditions and reset to naïve pluripotency (ERP006823, (Takashima et al., 2014)), conventional and chemically reset Shef6 cultures (ERP022538, (Guo et al., 2017)), epiblast-derived human naïve embryonic stem (HNES) cells (ERP014247 (Guo et al., 2016)) and independent cultures of clonal line of HNES1 (ERP022538, (Guo et al., 2017)), WIBR lines converted to a naïve state in 5i/L/A (SRP059227 (Ji et al., 2016)), WIS cells cultured in NSHM/4i (SRP045294 (Irie et al., 2015)), H1 and Lis1 lines independently cultured in NHSM/4i (SRP045911 (Sperber et al., 2015)), conventional H1 and Elf1 cells (SRP045911 (Sperber et al., 2015)) and extended pluripotent stem (EPS) cells reported to contribute to extraembryonic tissues (SRP074076 (Yang et al., 2017)). Reads were aligned to human genome build GRCh38/hg38 with STAR 2.5.2b (Dobin et al., 2013) using the two-pass method for novel splice detection (Engström et al., 2013). Read alignment was guided by GENCODE v25 (Harrow et al., 2012) human gene annotation from Ensembl release 87 (Yates et al., 2016) and splice junction donor/acceptor overlap settings were tailored to the read length of each dataset. Alignments to gene loci were quantified with htseq-count (Anders et al., 2015) based on annotation from Ensembl 87. Sequencing libraries with fewer than 500K mapped reads were excluded from subsequent analyses. Read distribution bias across gene bodies was computed as the ratio between the total reads spanning the 50th to the 100th percentile of gene length, and those between the first and 49th. Samples with ratio >2 were not considered further. Stage-specific outliers were screened by principal component analysis.

### Transcriptome analysis

Principal component and cluster analyses were performed based on log_2_ FPKM values computed with the Bioconductor packages *DESeq2* (Love et al., 2014), *Sincell* (Julia et al., 2015) or *FactoMineR* (Lê et al., 2008) in addition to custom scripts. T-distributed stochastic neighbor embedding (t-SNE, van der Maaten and Hinton, 2008) and diffusion maps were produced with the *Rtsne* (Krijthe, 2015) and *destiny* (Angerer et al., 2016) packages, respectively. Differential expression analysis was performed with *scde* (Kharchenko et al., 2014), which fits individual error models for the assessment of differential expression between sample groups. For global analyses, genes that registered zero counts in all single-cell samples in a given comparison were omitted. Euclidean distance and average agglomeration methods were used for cluster analyses. Fractional identity between preimplantation stages and *in vitro* cultured cells was determined via quadratic programming using the R package *DeconRNASeq* (Gong and Szustakowski, 2013). Average expression levels of cells comprising distinct stages were used as the “signature” dataset, and the relative identity of each culture protocol/sample group was computed by quadratic programming. Expression data are available in Supplemental Tables and through a web application to visualise transcription of individual genes in embryonic lineages (app.stemcells.cam.ac.uk/human-embryo).

### Selection of high-variability genes

Genes exhibiting the greatest expression variability (and thus contributing substantial discriminatory power) were identified by fitting a non-linear regression curve between average log_2_ FPKM and the square of coefficient of variation. Thresholds were applied along the x-axis (average log_2_ FPKM) and y-axis (log CV^2^) to identify the most variable genes. Depending on the samples compared, selection results were affected mostly by technical factors (library construction protocol, RNA-seq coverage, read distribution) where such properties differed between studies and datasets. To reduce these effects, when data from multiple studies were treated in a single analysis, we first identified variable genes independently for each dataset and then selected those genes common to each.

### Network analysis of biological processes

Statistical enrichment of Gene Ontology terms was computed with *GOstats* and DAVID 6.8 (Huang et al., 2009), using modulated genes as input (e.g. (between EPI and PrE cells, adjusted p<0.05). *Cytoscape* (Shannon et al., 2003) and the associated enrichment map plugin (Isserlin et al., 2014) were used for network construction and visualisation. For network diagrams, node size is scaled by the number of genes contributing to over-representation of biological processes; edges are plotted in width proportional to the overlap between gene sets. The ratio between up- and downregulated genes between cell populations (e.g. EPI and PrE) in each biological process is represented by colour shade (e.g. red, more genes upregulated in EPI; blue, more genes upregulated in PrE; see Fig. 4D).

### Evaluation of refined embryonic cell populations

To assess the accuracy of selected EPI, PrE and early ICM cells, we used the Weighted Gene Co-Expression Network Analysis unsupervised clustering method (WGCNA, Langfelder and Horvath, 2007) to identify specific modules of co-expressed genes in each developmental lineage. A soft power threshold of 10 was set to govern the correlation metric and a tree pruning approach (Langfelder et al., 2008) was implemented to merge similar modules (threshold 0.35). The minimum module size was set to 50 genes; from the modules computed, the top 50 genes with greatest intramodular connectivity were selected for subsequent co-expression network analysis.

### Identification of early and late preimplantation lineage markers

The R package *kohonen* was used to construct self-organizing transcriptome maps across embryonic stages. A matrix of 30×30 with hexagonal topology was used to map and identify areas of varying transcriptional activity. The *GOstats* package was used for stage-specific GO analyses considering genes with Z-score >1.5. Genes with Z-score <1.5 in all stages were used for the background set (universe). Annotation related to transcription factors, co-factors and chromatin remodelers was obtained from AnimalTFDB 2.0 (Zhang et al., 2015). Late lineage markers were selected as genes expressed in EPI or PrE cells with a transcriptional contribution more than 75% across all selected pre-implantation stages and minimum level of 10 FPKM. Early markers were identified as genes in later stages (from 8-cell morulae to either EPI or PrE lineages) with a transcriptional contribution of more than 75% across all selected pre-implantation stages. A fold change induction of at least four between lineages and minimum level of 10 FPKM in at least in one of the following stages was required: 8-cell, compacted morula, early ICM, EPI or PrE.

### Competing interests

GG and AS are inventors on a patent filing by the University of Cambridge relating to human naïve pluripotent stem cells.

### Funding

This work was supported by UK Biotechnology and Biological Sciences Research Council (BBSRC) research grant RG53615, UK Medical Research Council (MRC) programme grant G1001028, and institutional funding from the MRC and Wellcome Trust. AS is an MRC Professor.

## Supplementary Tables

**Table S1:** Differential expression analysis of Blakeley EPI, PrE and TE lineages.

**Table S2:** Differential expression analysis of EPI and PrE cells classified in this study.

**Table S3:** WGCNA of early ICM, EPI and PrE intramodular connectivity.

**Table S4:** Yan, Blakeley and Petropoulos samples with originally reported and reassessed lineage assignments.

**Table S5:** Normalised FPKM expression values for preimplantation stages.

**Table S6:** Stage-specific gene expression modules as defined by self-organising maps.

**Table S7:** Early and late lineage markers.

**Table S8:** Normalised averaged FPKM expression values for preimplantation stages and PSC cultures.

**Table S9:** Differential expression analysis of naïve versus conventional PSC.

## Supplementary Figures

**Fig. S1:**
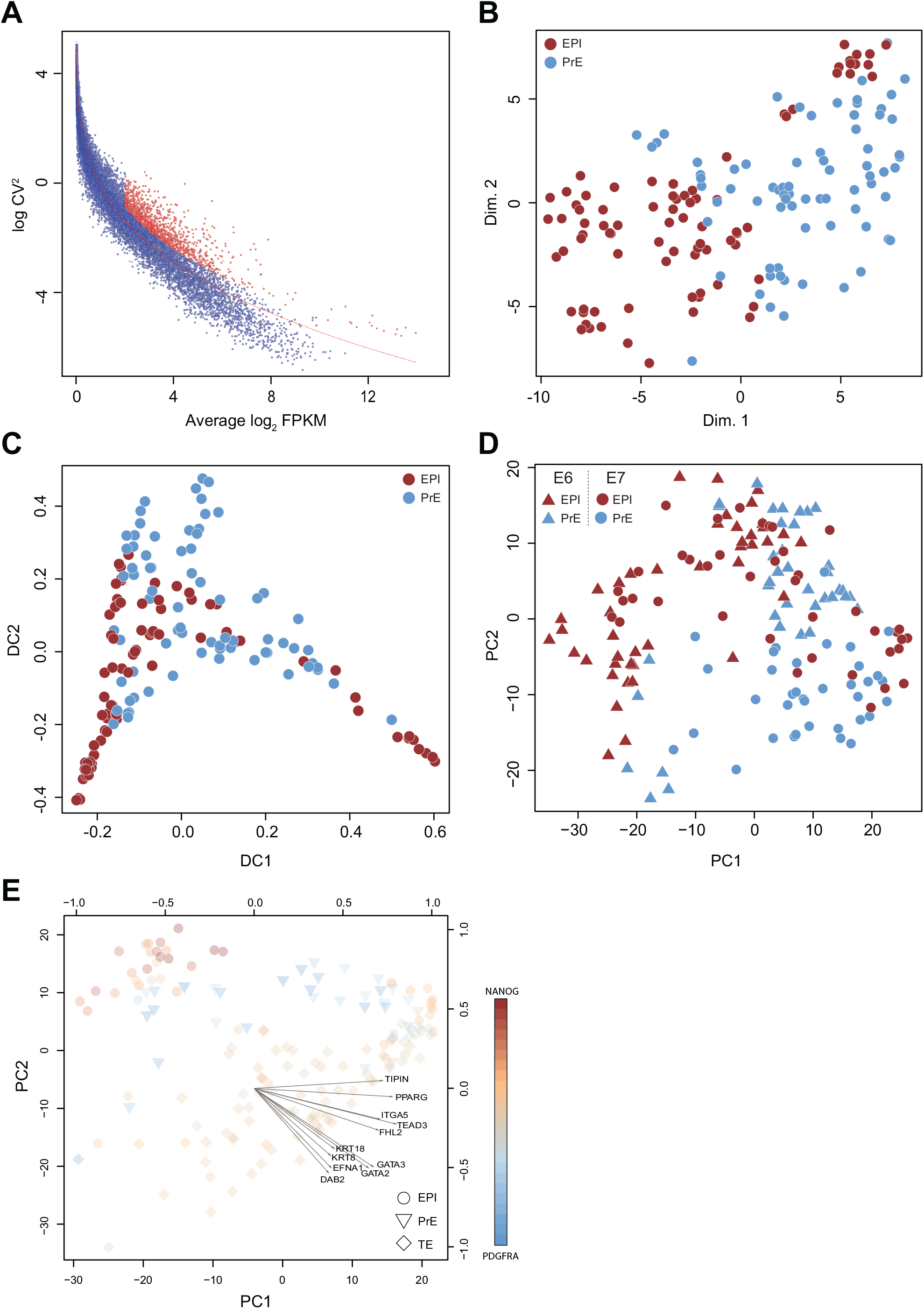
(A) Highly variable genes identified in E6 and E7 single-cell transcriptome data classified as EPI or PrE in Petropoulos et al. (2016). (B) t-SNE plot and (C) diffusion map of E6 and E7 EPI and PrE samples. (D) PCA of samples identified as EPI or PrE by Petropoulos et al. with developmental stage indicated. (E) Contribution of TE marker expression in Petropoulos immunosurgery samples.

**Fig. S2:**
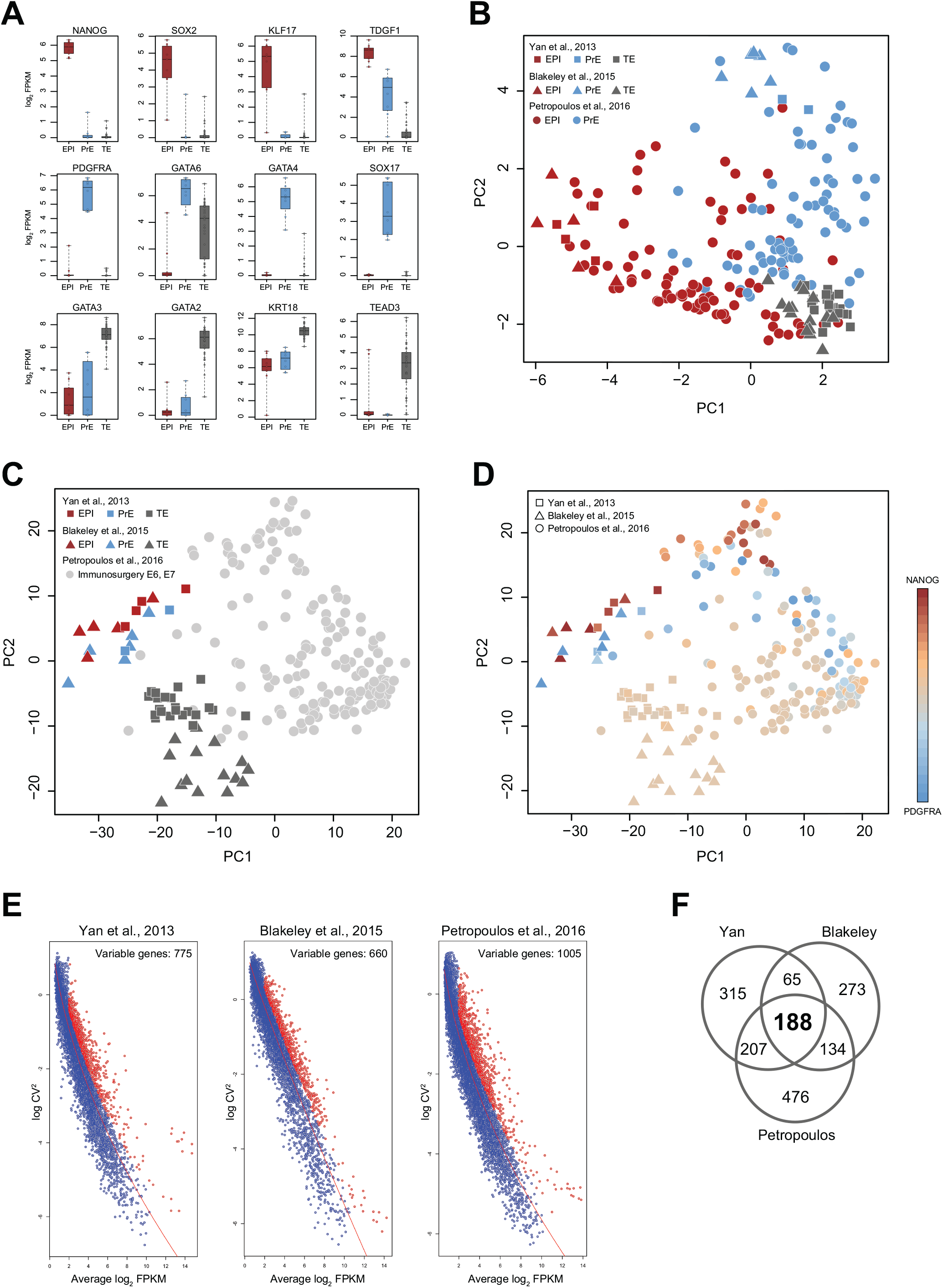
(A) Expression (log_2_ FPKM) of high-confidence lineage markers in cells profiled in Blakeley et al. (2015). (B) PCA of EPI, PrE and TE from Yan et al. (2013) and Blakeley datasets, together with E6 and E7 EPI and PrE samples from the Petropoulos study. (C) PCA based on variable genes (*n*=920) for EPI, PrE and TE samples from Yan and Blakeley datasets with the E6 and E7 immunosurgery subset from Petropoulos et al. (grey). (D) PCA as in C, where colours are scaled to the ratio of *NANOG* to *PDGFRA* expression. (E) Identification of variable genes for each dataset (log_2_ FPKM>2 and log CV^2^>0.5). (F) Intersection of variable genes common to Yan, Blakeley and Petropoulos samples.

**Fig. S3:**
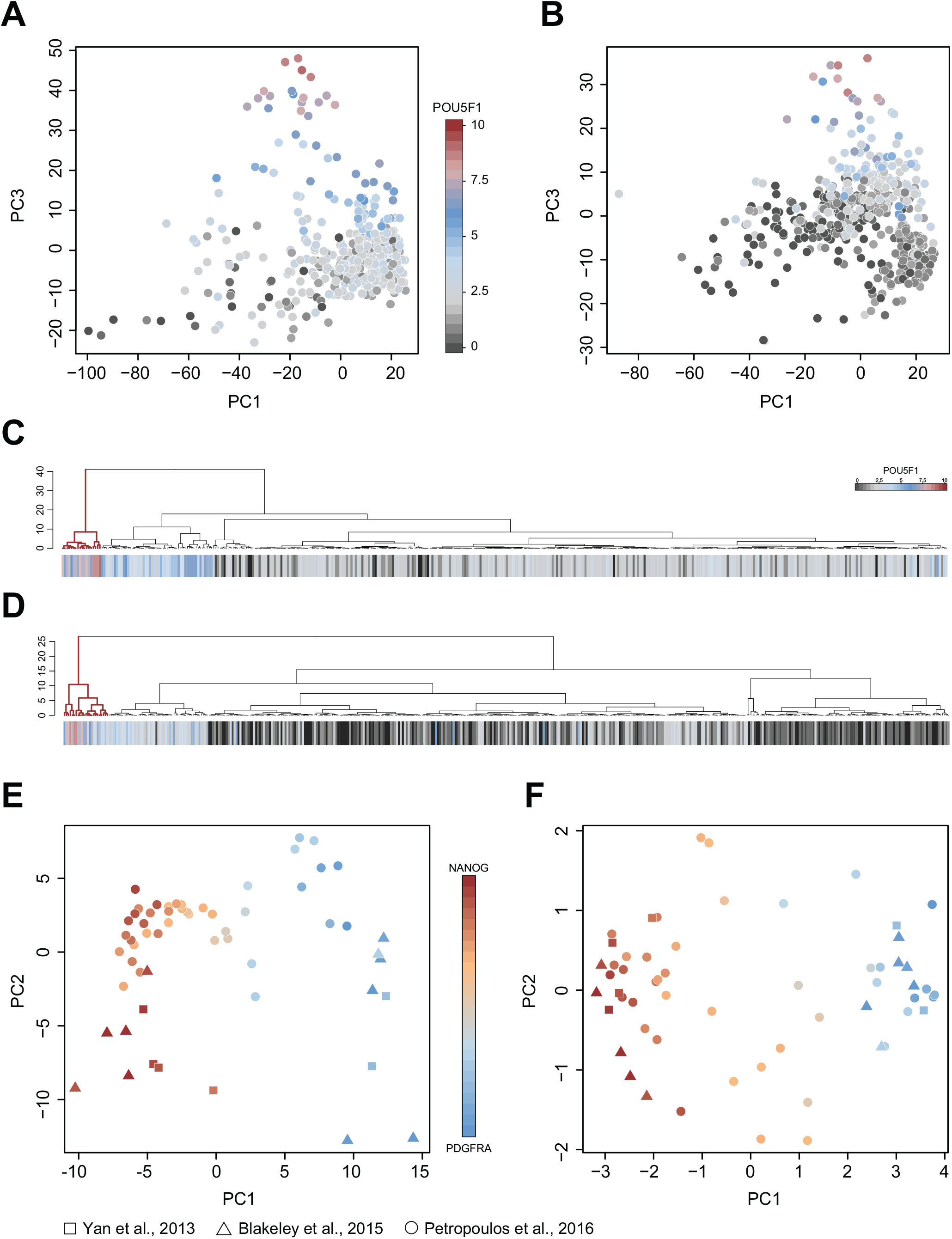
(A–B) PCA of E6 (A) and E7 (B) samples from Petropoulos et al. displaying the first and third principal components. Colours represent log_2_ *POU5F1* expression. (C–D) Dendrograms of E6 (C) and E7 (D) Petropoulos cells based on the third principal component in B, indicating *POU5F1* expression. *POU5F1*-high clusters are highlighted in red. (E) PCA based on common variable genes (*n*=181) of Petropoulos samples with high *POU5F1* levels as selected in C and D, together with EPI and PrE cells from Yan and Blakeley datasets. (F) PCA based on high-confidence markers as described in E.

**Fig. S4.**
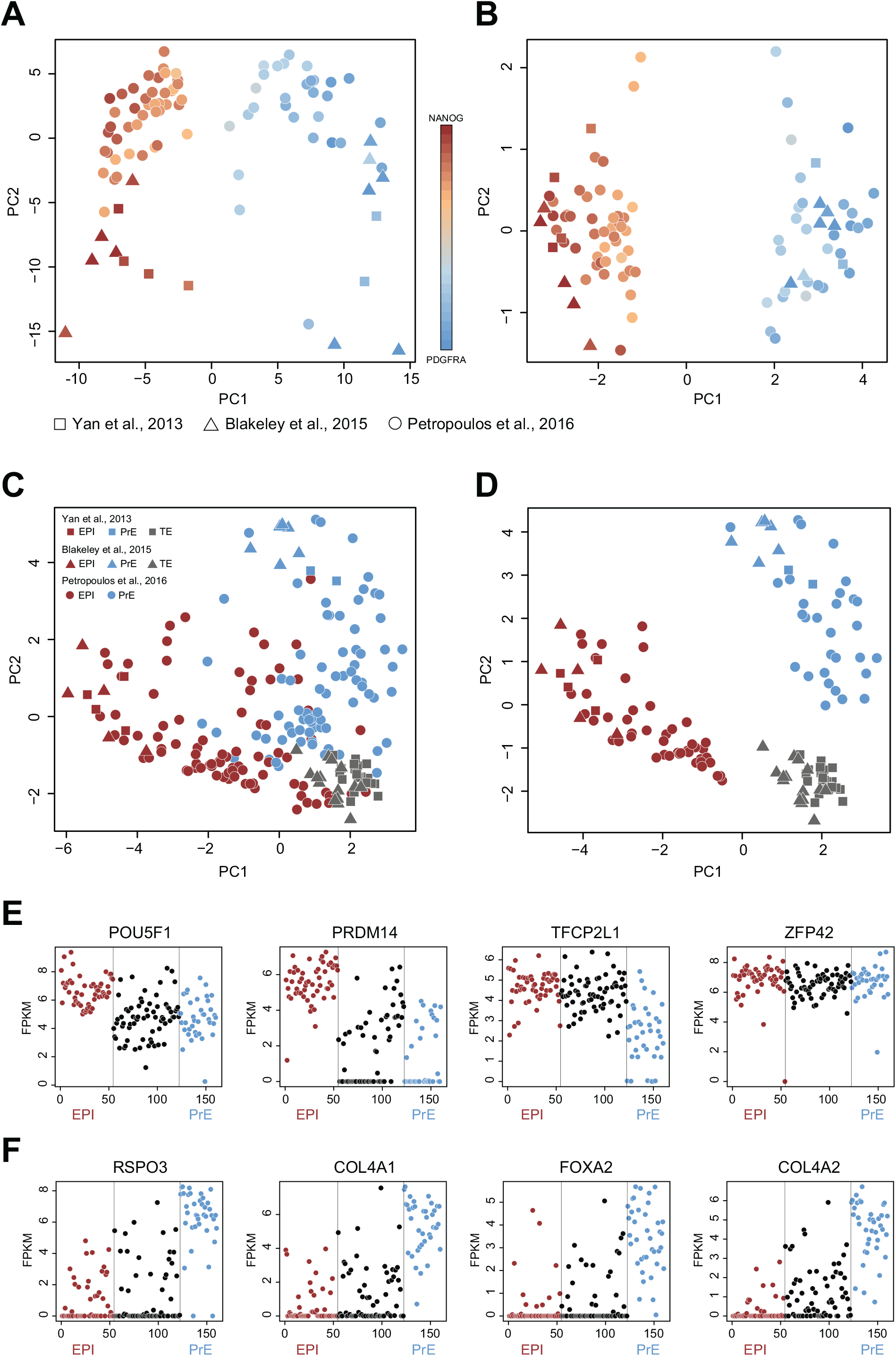
(A–B) PCA for selected EPI and PrE cells based on common variable genes (A, n=192) or lineage markers (B). Sample colours are scaled to the ratio of NANOG to PDGFRA expression. (C) PCA based on marker genes of E6 and E7 EPI and PrE cells as classified by Petropoulos et al. (red and blue circles), together with EPI, PrE and TE from Yan and Blakeley datasets. D) PCA based on marker genes of the EPI and PrE subselection in this study, together with EPI, PrE and TE from Yan and Blakeley datasets. (E, F) Single-cell dot plots of log_2_ FPKM values of the genes indicated. Cells are ordered along the x-axis to correspond with the dendrogram in Fig. 4A.

**Fig. S5:**
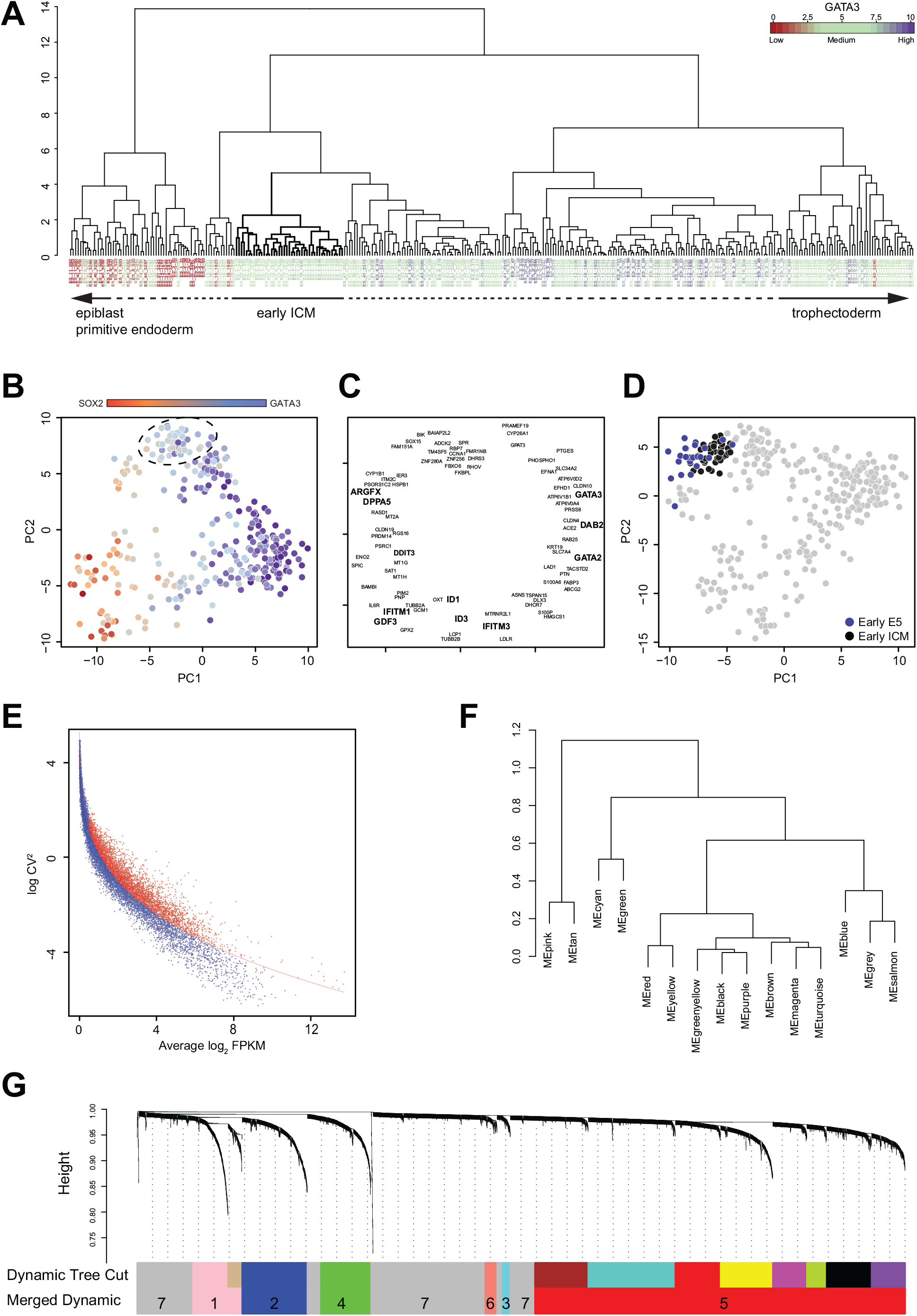
(A) Dendrogram of E5 cells from the Petropoulos dataset derived from the first two principal components of PCA based on highly variable genes (*n*=203). An early ICM population is highlighted in bold. Sample colours are proportional to *GATA3* expression. (B) PCA of E5 samples from Petropoulos et al. based on variable genes (*n*=205). Samples are scaled to the ratio of *SOX2* (red) versus *GATA3* (purple) levels. (C) Spatial positions of genes contributing to the principal components plotted in B. (D) PCA of E5 samples from Petropoulos et al. based on variable genes. Early ICM cells are indicated in black, early E5 cells as annotated by Petropoulos et al. are highlighted in blue. (E) Selection of variable genes (*n*=5097) for WGCNA analysis. (F) Hierarchical clustering of the initial 14 modules identified by WGCNA. (G) Co-expression of genes and modules before and after merging.

**Fig. S6:**
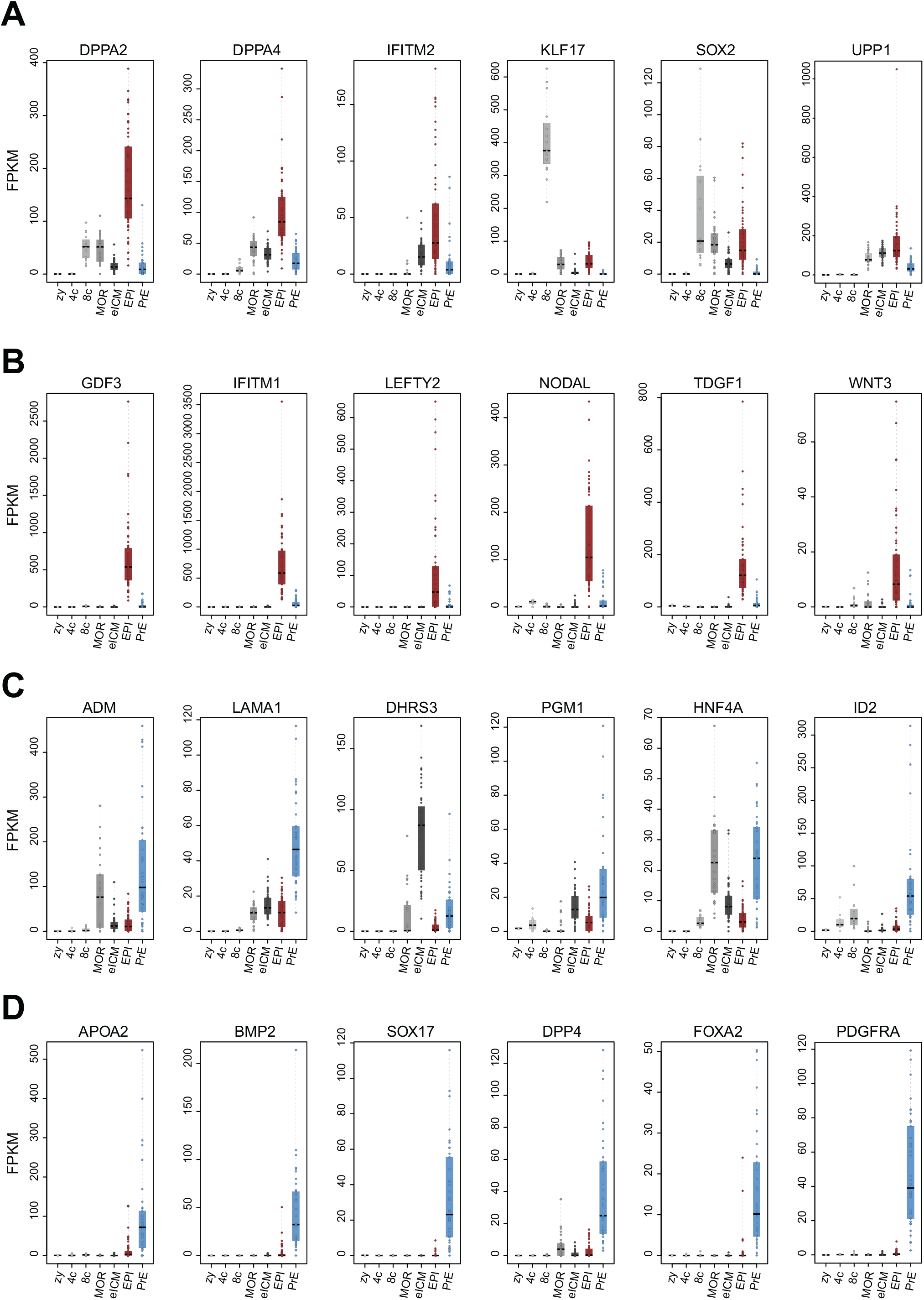
(A-D) Expression in FPKM of markers for (A) early EPI, (B) late EPI, (C) early PrE and (D) late PrE.

**Fig. S7:**
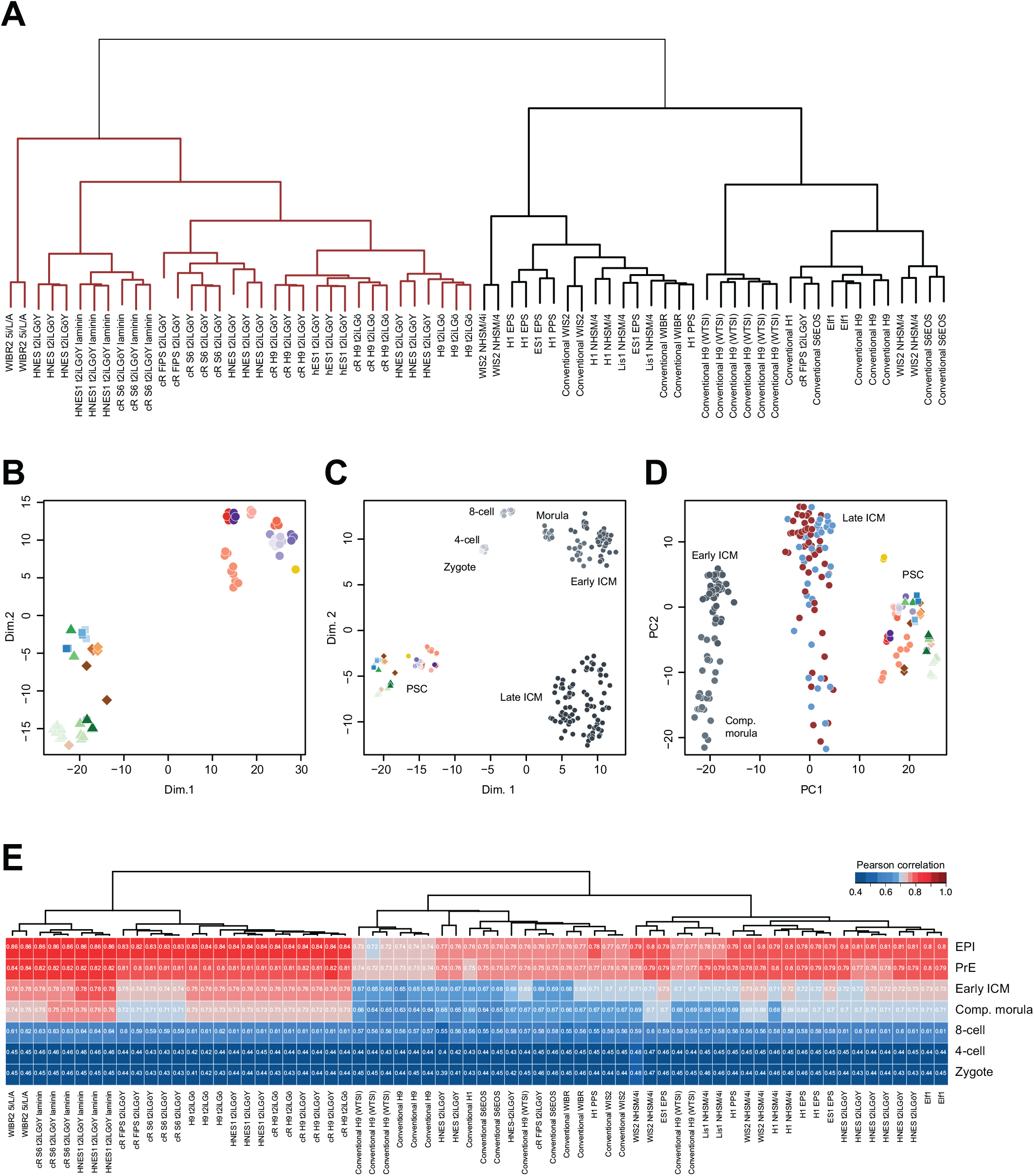
(A) Hierarchical clustering of in vitro PSC samples. (B) t-SNE of PSC based on genome-wide expression. (C) t-SNE as in (B) including all preimplantation embryo stages. (D) PCA based on highly variable genes (*n*=827) of PSC cultures together with compacted morula, early ICM and late ICM samples (red=EPI, blue=PrE). (D) Oneway clustering of Pearson correlation between averaged preimplantation stages and PSC cultures.

